# High-throughput pipeline for the *de novo* viral genome assembly and the identification of minority variants from Next-Generation Sequencing of residual diagnostic samples

**DOI:** 10.1101/035154

**Authors:** T Gallo Cassarino, D Frampton, R Sugar, E Charles, Z Kozlakidis, P Kellam

## Abstract

**Motivation:** The underlying genomic variation of a large number of pathogenic viruses can give rise to drug resistant mutations resulting in treatment failure. Next generation sequencing (NGS) enables the identification of viral quasi-species and the quantification of minority variants in clinical samples; therefore, it can be of direct benefit by detecting drug resistant mutations and devising optimal treatment strategies for individual patients.

**Results:** The ICONIC (InfeCtion respONse through vIrus genomiCs) project has developed an automated, portable and customisable high-throughput computational pipeline to assemble *de novo* whole viral genomes, either segmented or non-segmented, and quantify minority variants using residual diagnostic samples. The pipeline has been benchmarked on a dedicated High-Performance Computing cluster using paired-end reads from RSV and Influenza clinical samples. The median length of generated genomes was 96% for the RSV dataset and 100% for each Influenza segment. The analysis of each set lasted less than 12 hours; each sample took around 3 hours and required a maximum memory of 10 GB. The pipeline can be easily ported to a dedicated server or cluster through either an installation script or a docker image. As it enables the subtyping of viral samples and the detection of relevant drug resistance mutations within three days of sample collection, our pipeline could operate within existing clinical reporting time frames and potentially be used as a decision support tool towards more effective personalised patient treatments.

**Availability:** The software and its documentation are available from https://github.com/ICONIC-UCL/pipeline

**Supplementary information:** Supplementary data are available at *Briefings in Bioinformatics* online.

## 1 Introduction

Viruses are intracellular parasites and most are characterised by a high replication rate within their host. During replication the polymerase proteins are prone to transcription errors or mutations, with RNA viruses having the highest mutation rates. New genomes containing mutations are continuously generated and selected on the basis of their fitness to infect and to replicate within the host’s cells [1]. Average mutation rates of RNA viruses are about 10^−4^ – 10^−5^ errors per nucleotide copied or, based on average genomic sizes, about one mutation per genome copied[2]. Moreover, recombination of genomic parts increases the evolution dynamics and the genomic divergence within viral populations. One such example is the Human Immunodeficiency Virus (HIV), where the recombination is considered faster than the mutation rate[3]. These and other features suggest that a viral population is actually made of an ensemble of related mutants that can be described as a quasi-species and on which the selective pressure influences all the viruses as a single unit. Such high genomic variability allows the viral populations to survive challenges mounted by the host’s immune system and by antiviral agents, hindering the effective treatment of patients and making it difficult to eradicate infections[4]. Next-Generation Sequencing (NGS) coupled with bioinformatics analyses enables the high-throughput detection of genomic variants and the classification of known and novel viral species overcoming the need for expensive culturing and/or labour-intensive Sanger sequencing techniques[5].

Within the context of a clinical setting, NGS data can be applied on sequenced diagnostic samples to identify pathogens by assembling whole genomes and quantifying low-level drug resistance mutations below the 15–20% frequency sensitivity limit of the traditional Sanger sequencing technique[6].

Computational pipelines that employ NGS data have been developed recently to discover new pathogens[7, 8], to identify viral quasi-species[9, 10] and to detect genomic variants[11–14]. However, in order to impact patient treatment pathways, it is necessary for pipelines to be capable of combining a number of characteristics at the same time. These are the ability to be pathogen agnostic, to assemble *de novo* whole genomes, to report minority variants, to be scalable for high-throughput analyses, customisable and portable to different software environments.

To this end a computational pipeline for analysing NGS data was developed that meets all of these above requirements. The input data are unprocessed paired-end reads from residual clinical diagnostic samples obtained under appropriate ethics permissions, while the output reports the consensus genome and all minority variants present in the viral quasi-species. The pipeline is part of the ICONIC project, which uses viral genomic data to provide decision support towards the personalised treatment of patients; to guide hospital infection control responses; and to inform the surveillance and epidemiological responses to viral community outbreaks.

## 2 Methods

### 2.1 Pipeline Installation

The pipeline has been developed for analysing batches of sample paired-end reads on a High Performance Computing (HPC) cluster or on a server running the Son of Grid Engine scheduler (which must be installed separately). The software can be built either by the provided installation script or from a Dockerfile [https://www.docker.com/]. Both of these alternatives automatically: (1) download the pipeline dependencies, (2) configure the pipeline and (3) install it on the local appliance. Finally, the pipeline can be loaded as a bundle of Environment Modules[15] which can be used immediately. A pre-built binary version is also available as a docker image.

### 2.2 Read Analysis

The pipeline takes as input a batch of Illumina paired-end read FASTQ files generated from one or more samples, returning the consensus genome, minority variants and a set of statistics and QC metrics for each analysed sample and for the whole batch. It consists of a number of well-defined functional steps which allow the pipeline to be run from a particular starting point or to independently run a specific step. These functional steps can be optimised through a configuration file, which stores all the parameters used during the analysis. After parsing the input arguments, the pipeline initialises and sends an array job to the HPC cluster, which runs the analysis of each sample’s reads in parallel to facilitate rapid analysis. Each job is composed of six distinct steps as shown in Figure 1.

**Figure 1.**
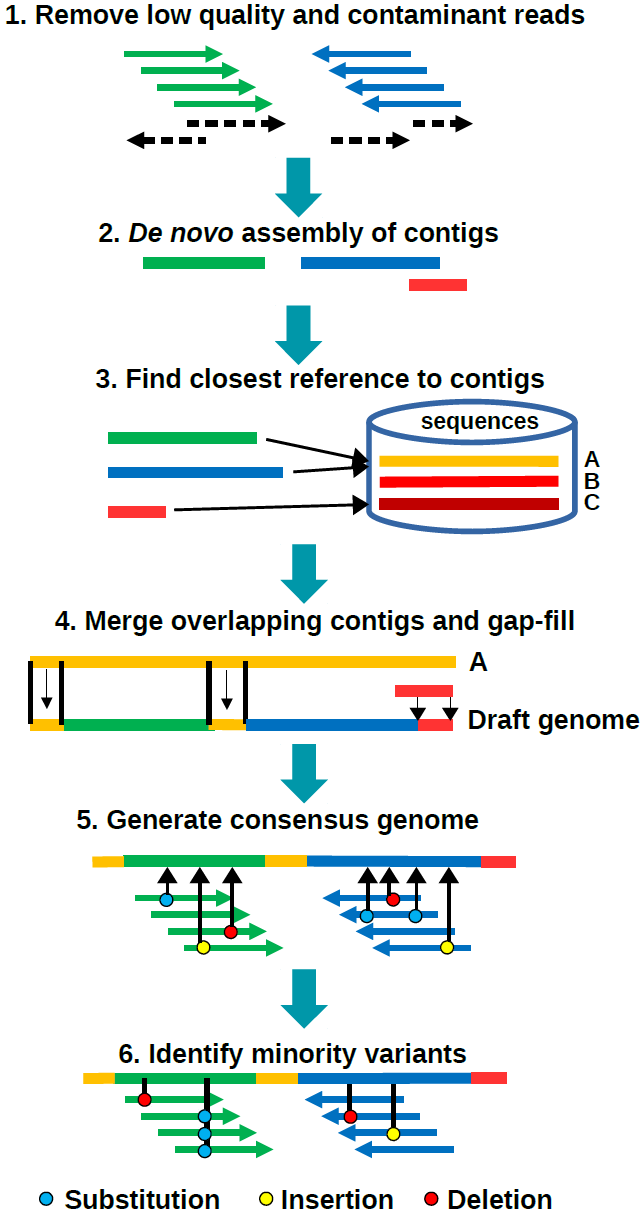
The workflow of the computational pipeline showing the major steps during the analysis of the sample reads.

The first workflow step takes as input the paired-end reads in either flat or compressed FASTQ format, keeping only high quality non-contaminant pairs. The two files are parsed through Trimmomatic16 (version 0.33) which removes or trims low quality reads (used with default settings, except for a sliding window Phred score cut-off of 30). The remaining read pairs are mapped against a user-defined decoy genome currently consisting of human chromosomes and an appropriate set of reference viral genomes, in order to remove unmapped reads and read pairs that map preferentially to the host than the pathogen. Mapping can be performed either with SMALT (version 0.7.6) (http://www.sanger.ac.uk/science/tools/smalt-0) or BWA[17] (version 0.7.12). Subsequently Samtools[18] (version 1.2) is used to manipulate and to extract data from the alignments and the read files. FastQC (version 0.11.3) (http://www.bioinformatics.babraham.ac.uk/projects/fastqc/) is run both before and after this filtering step to perform quality control checks and to spot potential sequencing biases. The results are saved in each sample’s output folder to allow subsequent inspection of read quality.

The second step generates contigs by *de novo* assembly of the filtered paired-end reads. In the current implementation, contigs are generated by the IVA *de novo* assembler[19] (version 1.0.0).

In the third step contigs are aligned with BLAST+[20] (version 2.2.30) against a local BLAST database of genomic sequences of the viral species specified as input, in order to identify the best match or closest reference. For segmented genomes (e.g. Influenza) this step identifies multiple best hits corresponding to each segment, thus allowing for potential reassortment.

The fourth step creates a draft genome from the assembled contigs. Using LASTZ21 (1.07.73) these are aligned to the previously found closest reference sequence or segments. Where contig alignments to the closest reference overlap, the draft sequence is taken from the contig with the highest read depth of coverage. Where contig alignments do not fully cover the coding sequence of the closest reference, gaps are filled with the missing sequence from the reference.

In the fifth step, an iterative approach is then used to generate the final consensus genome. Firstly, filtered reads are mapped to the draft genome (using SMALT or BWA) from which a pileup file is generated with Samtools. The pileup file is then parsed at each genomic position to quantify sequence variants relative to the draft genome. This variants data is analysed to identify positions where insertions or deletions are present in more than half of all mapped reads and/or positions where the most commonly reported mapped base differs from the draft genome. The draft genome is then corrected to account for these errors and the mapping and pileup steps are repeated until the genome converges to a stable sequence, i.e. when there are no more substitutions, insertions or deletions to be made. A maximum of 10 cycles are performed to avoid potentially infinite iterations.

In the sixth step, the reads are mapped for the last time to the consensus genome and the minority variants at each genomic position are identified using Samtools. To reduce false positive variant call, the pipeline allows user-defined cut-offs of variant read depth, variant frequency and consensus base frequency. Minority variants are also reported at variant frequency thresholds of 20% and 2%, in accordance with current Public Health England reporting practices and two additional bins of 10% and 5%.

At the end of the analysis, summary metrics and plots are generated for the consensus genome and for each genomic position; these metrics include genome length, read depth of coverage distribution, number of variants and strand bias. When all the jobs are completed, the pipeline collects the genome metrics of each sample and aggregates them to provide a set of statistics for the whole batch.

Optionally, it is possible to keep all intermediate files created during the analysis; however, this requires considerably more disk space.

The workflow also produces two log files during the execution of the analysis. The first stores all calls to external software (e.g. SMALT, Samtools, etc.) with associated memory usage and duration, while the second report the analysis steps: its verbosity level can be changed in the input options.

The pipeline save the final status of the sample analysis and the cause of any premature halt, (e.g. when *de novo* assembly fails due to an insufficient number of reads) thus assisting in identifying problematic samples.

Additional details on the input options and results can be found in the software documentation or in the GitHub repository. Moreover, new databases and decoy genomes can be added to the pipeline just by including their paths to the configuration file, thus allowing the pipeline to analyse NGS data of any virus.

### 2.3 Ethics permissions

The ICONIC study has REC approval (13/LO/1303) received on 20th August 2013, IRAS project ID 131373. The favourable opinion applies to all NHS sites taking part in the study, while additional permissions have been obtained from the NHS/HSC R&D offices of all partner sites prior to the start of the study.

## 3 Results

The capability of the pipeline to build *de novo* genomes was assessed using publicly available read datasets and, as an example, the results on a set of Human Respiratory Syncytial Virus (RSV) reads are shown. The pipeline performances were also tested on a bigger clinical sample datasets consisting of 341 Influenza virus samples sequenced at the Wellcome Trust Sanger Institute on an Illumina MiSeq machine.

### 3.1 Comparison with publicly available RSV genomes

A test batch of 39 publicly available sequences with deposited reads was selected at random for Human Respiratory Syncytial Virus (RSV), A and B subtypes, from the Sequence Read Archive (SRA). We created a local BLAST database of reference sequences from full genomes available in GenBank (as of 2015-10-01). With respect to the publicly available genomes, those built with the pipeline were longer in 19 samples (49%), shorter only in 7 (18%) due to the small number of reads remaining after the quality filtering step, and of comparable length in 13 samples (33%) (Details can be found in the Table_S1 of the supplementary material). The median length of built genomes was 96% of the corresponding reference. Therefore, our pipeline can build consistently full-size genomes and also contribute to the deposited public sequences by improving their genomic coverage.

### 3.2 Analysis of influenza clinical samples

The database used as reference set for the Influenza analysis was created using the Human Influenza full genomes belonging to any serotype downloaded from the NCBI’s Influenza Virus Resource (October 2015). The pipeline reported 169 genomes on a total of 342 (~50%) samples, while the rest did not contain enough viral reads to be *de novo* assembled into contigs. To overall assess the degree to which the consensus genomes covered the Influenza segments, these were aligned against a H1N1 sequence (strain: A/California/07/2009). Although the Influenza segments are relatively short (around 2000 bases for the longest one), the pipeline was able to create either partial or full-length segments (an example is shown by the genome depth in Figure S1 and Figure S2 for segment 1 and 4 respectively). The reads depth along the consensus genome qualitatively identified regions where the sequencing have been successful; Figure 2 illustrates one Influenza sample, showing that the number of aligned reads is higher at the ends of the segment than in the central region. Moreover, the number of soft-clipped reads highlighted regions where the alignment to the consensus genome is more difficult.

**Figure 2.**
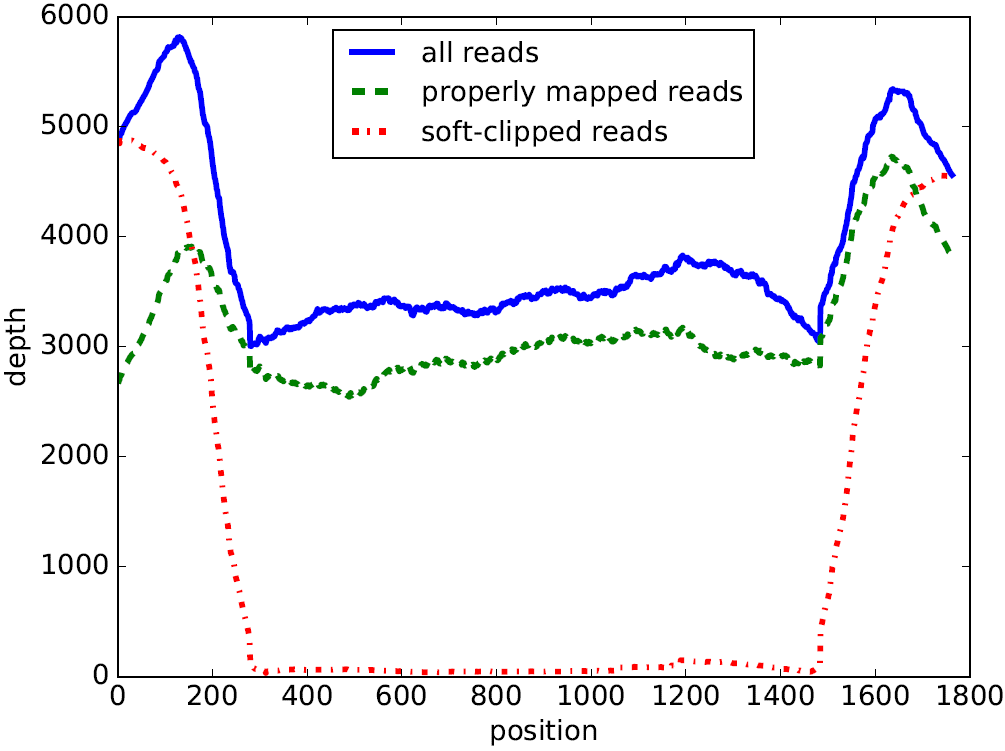
Read depth of coverage along the consensus sequence of segment 4 built for an Influenza sample. The blue line shows the number of aligned reads, the green dashed line shows the properly-mapped (according to the mapper) reads and the red dash-dotted line represents the number of soft-clipped reads.

For each consensus genome it was possible to inspect the minority variants in a table through: (1) their type (substitution, insertion or deletion), (2) the variant frequency and (3) the ratio between the amount of forward and reverse reads. By plotting the distribution of each single mutation against the consensus sequence (as shown in Figure 3), it was possible to identify genomic regions with high variability. The Influenza virus is characterized by a relatively low mutation rate; therefore, the number of minority variants at each position is quite small compared to high mutation rate viral genomes as HIV (as displayed for the Influenza segment 1 in Figure S3 in the Supplementary Material).

**Figure 3.**
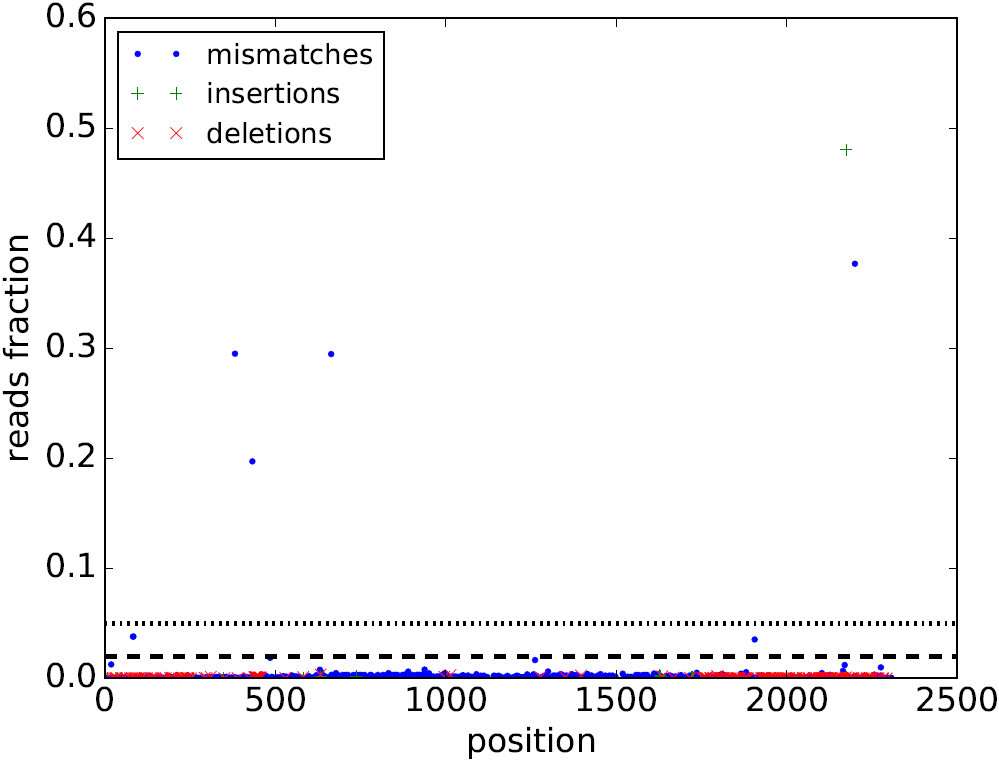
Single point mutations within the reads of an Influenza sample along the consensus genome. The blue dots show the mismatches, the green “plus” symbols show the insertions and the red crosses indicates the deletions, as the fraction of reads containing the mutation over the total number of reads aligned to the consensus genome.

The length of each assembled segment depends on the number of filtered reads available for that particular region, which can be assessed from the distribution of the median read depth (Figure S4 in the Supplementary Material). The segment coverage across the entire dataset is plotted in Figure S5 (Supplementary Material) through the distribution of the consensus genome coverage against its reference (Ns in the consensus genome are ignored); the median coverage was 100% for all segments and at least 50% for the first three segments in half of the dataset samples. The minority variant identified in the last step of the pipeline are aggregate from all the sample results and reported as well for each segment.

### 3.3. Performances

The analysis was performed using University College London’s HPC cluster “Legion” on 124 dedicated Dell C6220 nodes, where each node can work as a 16 core Symmetric Multi-Processing device with 64 GB of RAM. Legion runs an operating system based on Red Hat Enterprise Linux 7 with the Son of Grid Engine batch scheduler.

When the RSV samples were analysed, the pipeline ran for an average of about 2 hours per sample, and for less than 10 hours when the whole dataset. The analysis of the Influenza lasted on average less than 5 hours per sample and below 12 hours for the whole dataset (apart from 5 samples, for which the analysis took around 20 hours due to an exceptional long mapping time to the decoy genome).

The longest steps were the read alignment to the decoy genome and the *de novo* assembly, which were responsible for most of the pipeline execution time and increases with the nucleotide variability of the reads. All the software, except the scripts that build the consensus genome and manage the flow of the data through the workflow, was run in multi-threading mode using all the cores available to decrease the computational time of the analysis.

The memory used through the analysis had an initial peak around 10 GB, while aligning the raw reads to the decoy genome, and it reached 2 GB during the *de novo* assembly step; otherwise, the average memory load is about 500 MB.

Finally, Docker images have a negligible impact on the performances[22] and this holds true especially for our pipeline, since it is contained in a single Docker image.

### 3.4 Portability

The pipeline can be installed either from a Dockerfile or by an installation script on a GNU/Linux system, or it can be ran directly as a docker image. The former installation method can be used on a dedicated single-user appliance, while the latter is better suited for environments shared among multiple users, hence addressing different set up needs. Often Docker cannot be installed on academic clusters for security reasons, as the ability of running docker images is equivalent to having access to root privileges [Docker Security: https://docs.docker.com/engine/articles/security/]. In these cases Environmental Modules are used to make sure all software dependencies are loaded with the correct version. This option was necessary to address the between-institutions portability.

The pipeline and its documentation can be downloaded or cloned from the GitHub repository at https://github.com/ICONIC-UCL/pipeline

## 4 Discussion

Motivated by the increasing potential to apply NGS on clinical viral samples as a method to improve the treatment of patients affected by viral diseases, a high-throughput computational pipeline was developed as part of the ICONIC project to assemble viral consensus genomes *de novo* and to detect minority variants in viral residual clinical samples. As the pipeline accepts raw reads, it is possible to analyse sequencing data as soon as they are produced without the need of any pre-processing step. Moreover, the pipeline can analyse reads from any viral genome, segmented or not, for which at least one sequence, even partial, already exists. The consensus genome and the associated minority variants can be used to identify the dominant viral subtype and to quantify the variants occurring at specific positions and identifying the presence of drug resistance mutations. Furthermore, the possibility to run the pipeline in high-throughput mode is essential to facilitate analyses of potentially large numbers of patient samples during a seasonal outbreak of conditions of viral origin. Given the variety of information technologies employed within different clinical environments, the pipeline was designed to be easily ported to any cluster or server running a GNU/Linux system.

A set of publicly available RSV sample sequences were compared to the genomes generated by the pipeline presented here, starting from the deposited reads. Full genomes were generated in most of the samples, with half giving longer alignments to reference than those that are currently publicly available. In this way, the pipeline can successfully and efficiently assemble *de novo* viral genomes and could potentially be used to replace and update the data deposited on public archives. The subsequent analyses of Influenza reads on an HPC cluster, sequenced from clinical residual samples, confirmed the above results. The analysis lasted only a few hours and requiring an operationally reasonable amount of memory for each sample, thus capable of processing batches of hundreds samples overnight on a typical HPC cluster. Such a timeframe, coupled with the easy portability, allows the pipeline to be suitable to high pressure settings, such as clinical settings, where reporting turnaround times are compressed, especially in the case of diseases of viral aetiology.

It is not difficult to imagine that using current capability, a diagnostic report could be provided to a clinician within an actionable time window, as in our experience the end-to-end process from patient sampling to genome assembly and clustering reporting takes approximately five days. One of the main strengths of the pipeline is that it can be utilised on sequencing data of any known virus, to generate *de novo* full length viral genomes, in which the sample quality is very variable and the virus subtype is unknown. These features empower our software to be eventually deployed in clinical settings as a decision support tool towards a personalised patient treatment and the improved information management of hospital infections.

## Acknowledgments

We would like to thank Chiara Garattini (Intel Corporation) for the useful suggestions and discussions during the preparation of the manuscript and Christophe Fraser, Chris Wymant and Oliver Ratmann from the BEEHIVE consortium for their input during the pipeline’s initial development and the PANGEA_HIV consortium for facilitating these discussions.

## Key points

The ICONIC high-throughput computational pipeline *de novo* assembles viral genomes and quantifies minority variants.
It uses Illumina paired-end reads sequenced from residual diagnostic samples.
It could operate within existing clinical reporting time frames and potentially be used as a decision support tool towards more effective personalised patient treatments.

## Legends for the supplementary material

Table S1. Sample identifiers from the Sequence Read Archive with the length of the associated deposited RSV genome and of the consensus sequence built by the ICONIC pipeline.

Figure S1. Genome depth of coverage of the segment 1 within the Influenza batch. The blue line shows the number of genomes aligned at each position of the reference sequence.

Figure S2. Genome depth of coverage of the segment 4 within the Influenza batch. The blue line shows the number of genomes aligned at each position of the reference sequence.

Figure S3. Minority variants of the segment 1 within the Influenza sample dataset. Each dot represents the absolute number of minority variants found in the sample reads aligned to the consensus genome.

Figure S4. Median read depth of coverage for each segment across the Influenza batch. Median is shown as red line, 25 and 75 QRT as box, 95 QRT as whiskers and outliers as plus signs.

Figure S5. Segment coverage from all the sample in the Influenza batch. All segments have median equal to 100%, segments 4 and 5 have only the 95 QRT below 100%, while segments 7 and 8 have all coverages at 100%. Median is shown as red line, 25 and 75 QRT as box, 95 QRT as whiskers and outliers as plus signs.

## References

1. Moya A, Holmes EC, Gonzalez-Candelas F. The population genetics and evolutionary epidemiology of RNA viruses, Nat Rev Microbiol 2004;2:279–288.

2. Sanjuan R, Nebot MR, Chirico N et al. Viral mutation rates, J Virol 2010;84:9733–9748.

3. Rhodes T, Wargo H, Hu WS. High rates of human immunodeficiency virus type 1 recombination: near-random segregation of markers one kilobase apart in one round of viral replication, J Virol 2003;77:11193–11200.

4. Domingo E, Menendez-Arias L, Quinones-Mateu ME et al. Viral quasispecies and the problem of vaccine-escape and drug-resistant mutants, Prog Drug Res 1997;48:99–128.

5. Quinones-Mateu ME, Avila S, Reyes-Teran G et al. Deep sequencing: becoming a critical tool in clinical virology, J Clin Virol 2014;61:9–19.

6. Tsiatis AC, Norris-Kirby A, Rich RG et al. Comparison of Sanger sequencing, pyrosequencing, and melting curve analysis for the detection of KRAS mutations: diagnostic and clinical implications, J Mol Diagn 2010;12:425–432.

7. Naccache SN, Federman S, Veeraraghavan N et al. A cloud-compatible bioinformatics pipeline for ultrarapid pathogen identification from next-generation sequencing of clinical samples, Genome Res 2014;24:1180–1192.

8. Zhao G, Krishnamurthy S, Cai Z et al. Identification of novel viruses using VirusHunter––an automated data analysis pipeline, PLoS One 2013;8:e78470.

9. Prosperi MC, Salemi M. QuRe: software for viral quasispecies reconstruction from next-generation sequencing data, Bioinformatics 2012;28:132–133.

10. Topfer A, Marschall T, Bull RA et al. Viral quasispecies assembly via maximal clique enumeration, PLoS Comput Biol 2014;10:e1003515.

11. Wilm A, Aw PP, Bertrand D et al. LoFreq: a sequence-quality aware, ultra-sensitive variant caller for uncovering cell-population heterogeneity from high-throughput sequencing datasets, Nucleic Acids Res 2012;40:11189–11201.

12. Watson SJ, Welkers MR, Depledge DP et al. Viral population analysis and minority-variant detection using short read next-generation sequencing, Philos Trans R Soc Lond B Biol Sci 2013;368:20120205.

13. McElroy K, Zagordi O, Bull R et al. Accurate single nucleotide variant detection in viral populations by combining probabilistic clustering with a statistical test of strand bias, BMC Genomics 2013;14:501.

14. Verbist BM, Thys K, Reumers J et al. VirVarSeq: a low-frequency virus variant detection pipeline for Illumina sequencing using adaptive base-calling accuracy filtering, Bioinformatics 2015;31:94–101.

15. Furlani JL, Osel PW. Abstract Yourself With Modules. Proceedings of the 10th USENIX conference on System administration. Chicago, IL: USENIX Association, 1996, 193–204.

16. Bolger AM, Lohse M, Usadel B. Trimmomatic: a flexible trimmer for Illumina sequence data, Bioinformatics 2014;30:2114–2120.

17. Li H. Aligning sequence reads, clone sequences and assembly contigs with BWA-MEM. ArXiv e-prints. 2013.

18. Li H, Handsaker B, Wysoker A et al. The Sequence Alignment/Map format and SAMtools, Bioinformatics 2009;25:2078–2079.

19. Hunt M, Gall A, Ong SH et al. IVA: accurate de novo assembly of RNA virus genomes, Bioinformatics 2015;31:2374–2376.

20. Camacho C, Coulouris G, Avagyan V et al. BLAST+: architecture and applications, BMC Bioinformatics 2009;10:421.

21. Harris RS. Improved pairwise alignment of genomic DNA. College of Engineering. The Pennsylvania State University, 2007.

22. Di Tommaso P, Palumbo E, Chatzou M et al. The impact of Docker containers on the performance of genomic pipelines, PeerJ 2015;3:e1273.

